# Multimodal illumination platform for 3D single-molecule super-resolution imaging throughout mammalian cells

**DOI:** 10.1101/2024.02.08.579549

**Authors:** Tyler Nelson, Sofía Vargas-Hernández, Margareth Freire, Siyang Cheng, Anna-Karin Gustavsson

**Author notes:** These authors contributed equally to this work (Tyler Nelson, Sofía Vargas-Hernández).

## Abstract

Single-molecule super-resolution imaging is instrumental for investigating cellular architecture and organization at the nanoscale. Achieving precise 3D nanometric localization when imaging structures throughout mammalian cells, which can be multiple microns thick, requires careful selection of the illumination scheme in order to optimize the fluorescence signal to background ratio (SBR). Thus, an optical platform that combines different wide-field illumination schemes for target-specific SBR optimization would facilitate more precise, 3D nanoscale studies of a wide range of cellular structures. Here we demonstrate a versatile multimodal illumination platform that integrates the sectioning and background reduction capabilities of light sheet illumination with homogeneous, flat-field epi-and TIRF illumination. Using primarily commercially available parts, we combine the fast and convenient switching between illumination modalities with point spread function engineering to enable 3D singlemolecule super-resolution imaging throughout mammalian cells. For targets directly at the coverslip, the homogenous intensity profile and excellent sectioning of our flat-field TIRF illumination scheme improves single-molecule data quality by providing low fluorescence background and uniform fluorophore blinking kinetics, fluorescence signal, and localization precision across the entire field of view. The increased contrast achieved with LS illumination, when compared with epi-illumination, makes this illumination modality an excellent alternative when imaging targets that extend throughout the cell. We validate our microscopy platform for improved 3D super-resolution imaging by two-color imaging of paxillin – a protein located in the focal adhesion complex – and actin in human osteosarcoma cells.

## 1. Introduction

Fluorescence single-molecule localization microscopy (SMLM) [1–5] is playing a crucial role in addressing fundamental questions in molecular and cellular biology [6–10]. In particular, single-molecule super-resolution imaging (SRI) has made it possible to elucidate previously unknown properties of various cellular structures, at length scales spanning protein assemblies and macromolecular complexes [11,12] to cellular architecture and organization studies in three dimensions (3D) [13–15]. Achieving precise 3D nanometric localization for any given sample within such a diverse range of structural targets requires optimization of the fluorescence signal to background ratio (SBR) to minimize the localization uncertainty [16,17]. When imaging throughout mammalian cells, which can be many microns thick, the SBR should thus be optimized with the region of study in mind [18–24].

One approach to maximize the SBR and improve the localization precision is to reduce the out-of-focus fluorescence background by carefully selecting the illumination method [8,17]. However, there is often a tradeoff between optimal performance and simplicity of implementation for illumination schemes. Wide-field epi-illumination is a commonly used method for SMLM throughout a cell, but since the entire sample is illuminated, major drawbacks include high fluorescence background and increased risk of photobleaching and photodamage. Confocal illumination offers significant background reduction with the use of a pinhole to reject light originating from out-of-focus planes at the expense of relatively slow acquisition speeds, given its raster scanning nature. The speed can be improved by spinning disk implementations [25,26], but confocal approaches require higher peak illumination intensities to achieve the same signal and they suffer from increased risk of photobleaching and photodamage as the beam illuminates the entire depth of the sample [27]. A wide-field alternative that offers exquisite optical sectioning for reduced fluorescence background, photobleaching, and photodamage is total internal reflection fluorescence (TIRF) illumination. TIRF is easy to implement and has been used extensively for SMLM applications, but its use is restricted to a few hundred nanometers above the coverslip [28], making it incompatible with imaging throughout thick samples. An alternative to circumvent such limitations is light sheet (LS) illumination [29–31]. LS illumination is a wide-field technique where the sample is illuminated with a plane of light introduced orthogonally to the detection axis at the image plane. LS illumination is well-suited for imaging thick mammalian cells since it can provide excellent optical sectioning throughout the entire sample [32,33].

Another important consideration for precise 3D SMLM in large samples, such as mammalian cells, is the homogeneity of the localization precision and the number of localizations throughout the field-of-view (FOV). Generally, lasers with Gaussian beam profiles are used for illumination in SMLM. This inhomogeneous intensity profile results in decreased signal towards the edges of the FOV, which effectively degrades the localization precision in those areas [34,35]. Also, heterogenous illumination across the sample causes high variability in photobleaching rates [35,36] and other photophysical behaviors of fluorophores [34,35], complicating the extraction of quantitative information from the singlemolecule data. Overcoming these limitations has motivated the implementation of top-hat, or flat-field (FF), illumination profiles for SMLM, which have been achieved using multiple different approaches, including the fabrication of waveguide chips [37,38], the use of diffractive [35] and refractive [36] beam-shapers, microlens arrays [39], galvanometric scanning mirrors [34], and multimode fibers [40–42]. Each of these implementations has inherent tradeoffs in cost, complexity of setup, and adaptability for multimodal illumination. Nonetheless, it has been extensively demonstrated that the use FF profiles for wide-field epiand TIRF illumination significantly improves the quality of SMLM data [34–36] by the homogenization of localization precision, photobleaching rates, and blinking kinetics throughout the entire FOV.

Consequently, an optical platform that integrates different wide-field illumination schemes for SBR optimization with homogeneous illumination and 3D SRI capabilities would facilitate more precise, 3D nanoscale studies of a wide range of cellular structures. Here we demonstrate a flexible multimodal platform for 3D single-molecule SRI that integrates the optical sectioning capabilities of a tilted LS with FF epi- and FF TIRF illumination. A two-channel 4f system enables point spread function (PSF) engineering for 3D SRI. Fast switching between illumination modalities is enabled by a galvanometric mirror, and the alignment and steering of the LS is completely decoupled from the optical path of the FF epi- and FF TIRF illumination setup. Our design is cost-efficient, made from primarily commercially available parts, is easy to align, with a freely available CAD of the platform to facilitate construction and implementation. We demonstrate the performance of our platform for 3D SRI in thick samples by imaging of paxillin and the actin cytoskeleton in human osteosarcoma (U-2 OS) cells.

## 2. Methodology

Details of sample preparation, imaging conditions, and data analysis can be found in the supplemental document.

A schematic of the optical platform is shown in Fig. 1a (see supplemental Fig. S1 for CAD rendering of the setup and supplemental CAD file for the full design of the setup and for all custom parts).

**Fig. 1.**
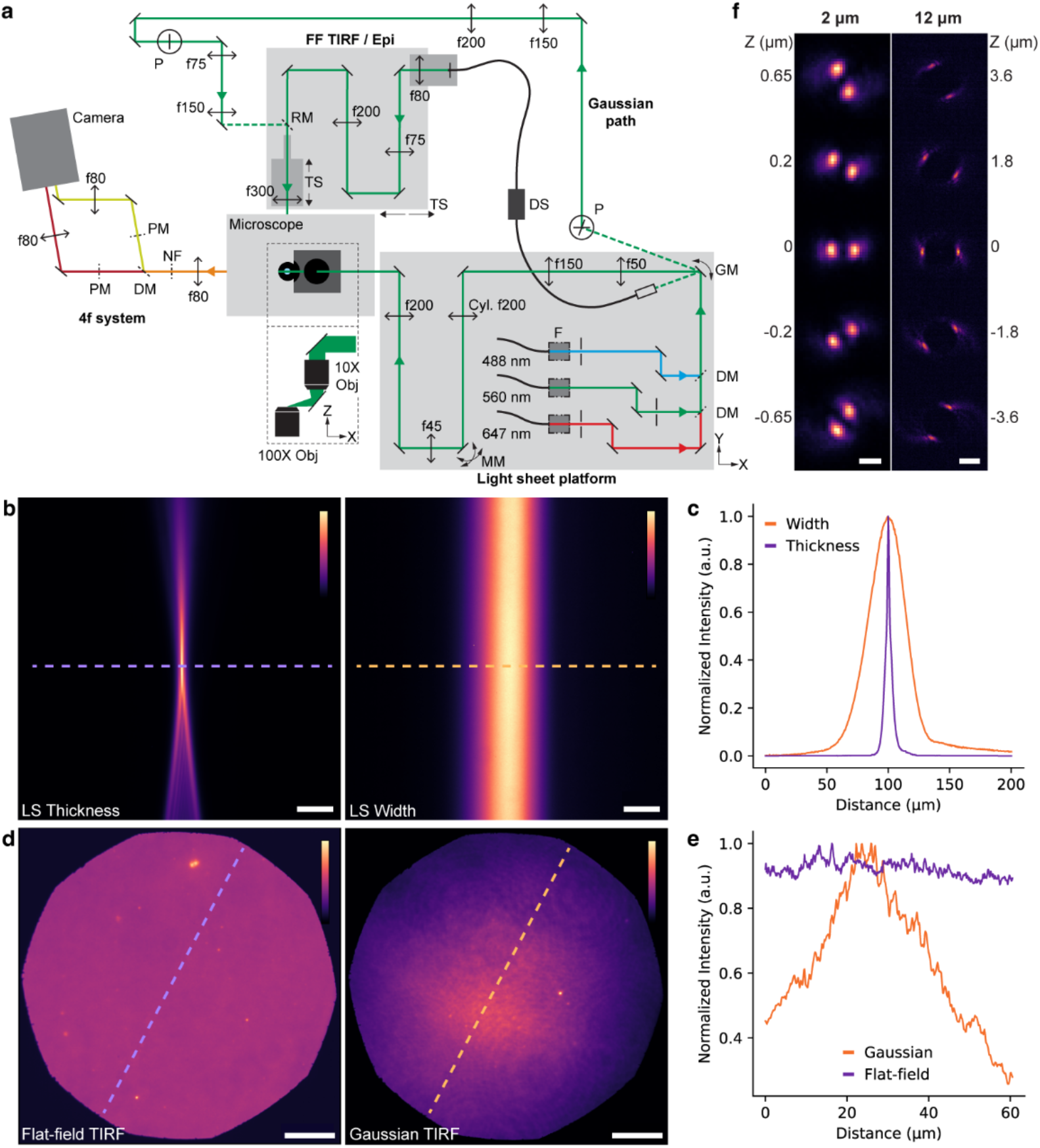
Design and performance of the multimodal illumination platform. (a) Simplified schematic of the microscopy platform (not to scale). Definitions of optical elements abbreviations: cylindrical lens (Cyl.), despeckler (DS), dichroic mirror (DM), galvanometric mirror (GM), motorized mirror (MM), notch filter (NF), objective lens (Obj), periscope (P), phase mask (PM), removable mirror (RM), translation stage (TS). f indicates lenses with the indicated focal lengths in mm. (b) Images of the light sheet (LS) in fluorescent solution showing the LS thickness (left) and width (right). Scale bars are 10 μm. (c) Graph with line scans of the LS images shown in (b) at the indicated dashed lines used to determine the LS thickness and width. (d) Images of a dense layer of fluorescent beads showing the flat-field (FF, left) and Gaussian (right) TIRF illumination intensity profiles. Scale bars are 10 μm. (e) Graph with line scans of the images shown in (d) at the indicated dashed lines used to quantify the uniformity of the FF intensity profile compared to the Gaussian intensity profile. (f) Images of fluorescent beads at the indicated axial positions demonstrating the double-helix PSFs with 2 μm (left) and 12 μm (right) axial range. Scale bars are 1 μm and 3 μm for left and right panels, respectively. Color bars indicate the linear color scale used. Each image’s contrast is normalized independently.

Three illumination lasers (488 nm, 200 mW, Coherent; 560 nm, 1000 mW and 647 nm, 1000 mW, both MPB Communications; all continuous wave (CW)) were mounted on an elevated 12×18” breadboard (MB1218, Thorlabs) positioned to the side of an inverted microscope (IX-83, Olympus) with a high NA detection objective (UPLAPO100XOHR, 100X, NA 1.5, Olympus) and XYZ stages (OPH-XYS-O and OPH-PINANO-XYZ both Physik Instrumente) for sample movement. Each laser was spectrally filtered (FF01-472/27-25, FF01-554/23-25, FF01-631/36-25, respectively; all Semrock), and circularly polarized (linear polarizers: LPVISA050-MP2 (488 nm) and LPVISC050-MP2 (560 nm and 647 nm), all Thorlabs; and quarter-wave plates: Z-10-A-.250-B-488, Z-10-A-.250-B-556, and Z-10-A-.250-B-647, respectively, all Tower Optics). Individual shutters (see supplemental Fig. S1 for CAD rendering and supplemental CAD file for the 3D printed mounts) were used to toggle the lasers (VS14S2T1 with VMM-D3 three-channel driver, Uniblitz, Vincent Associates). The beams were then merged into a single beam path using dichroic mirrors (DMLP505T and DMLP567T, both Thorlabs) and directed to a galvanometric mirror (GVS211, ±20° angular range, Thorlabs). The galvanometric mirror was used to direct the beam into either the LS (0°), FF (+20°), or Gaussian (−20°) path. The shutters and galvanometric mirror were connected to a computer via a multifunction I/O device (PCIe-6353, National Instruments) and BNC rack-mount connector (BNC-2090A, National Instruments) to enable automatic control.

The LS path was designed to fit entirely on the elevated breadboard. After the galvanometric mirror, the beam for the LS path was expanded and collimated using a lens telescope (f = 50 mm, AC254-050-A, Thorlabs and f = 150 mm, AC254-150-A, Thorlabs). Next, the beam was focused in one dimension by a cylindrical lens (f = 200 mm, LJ1653RM-A, Thorlabs) in a rotating mount (RSP1, Thorlabs) onto a motorized mirror (8807, Newport) positioned one focal length from the cylindrical lens. Two lenses (f = 45 mm, AC254-045-A, Thorlabs and f = 200 mm, ACT508-200-A, Thorlabs) were placed in 4f configuration to image the plane of the motorized mirror onto the back focal plane of a long working distance illumination objective (MY10X-803, 10X, NA 0.28, Mitutoyo). In order to direct the beam vertically downward through the illumination objective, a mirror in a tip/tilt mount was placed in a cage assembly affixed to the breadboard and mounted directly on top of the illumination objective mount (ST1XY-S, Thorlabs). Next, a mirror was glued in a custom 3D printed mount (see supplemental Fig. S1 and supplemental CAD for design) attached to the illumination objective to send the LS beam at an 11° angle relative the horizontal plane into the sample. To allow easy movement and alignment of the LS throughout the sample, the breadboard was placed on an XY stage (401, Newport) which was mounted on top of a large Z translational stage (281, Newport).

To achieve FF TIRF/epi-illumination, the galvanometric mirror was set at +20° to direct the beam via a fiber collimator (64786, NA 0.55, Edmund Optics) in a tip/tilt mount into a 200 μm square-core multi-mode fiber with an incorporated de-speckler for further beam homogenization (F-DS-ASQR200-SMA, Newport). To facilitate easy switching between epi- and TIRF illumination, the end of the multi-mode fiber and all subsequent optics in the FF path were positioned on a single linear translation stage (TBB1212, Thorlabs), mounted to the optical table at the rear of the microscope. Thus, translating the beam at the back focal plane of the microscope objective (UPLAPO100XOHR, 100X, NA 1.5, Olympus) to switch between epi-illumination and TIRF only requires moving the translational stage position without further optics alignment. A 4×6” breadboard was appended to one side of the translational stage to provide more space to mount the multi-mode fiber output and a collimating lens (f = 80 mm, AC508-080-A, Thorlabs). To adjust the size of the beam, a lens telescope (f = 75 mm, AC508-075-A, Thorlabs and f = 200 mm, 45417, Edmund Optics, respectively) in 4f-configuration was placed parfocally to a Köhler lens (f = 300 mm, 45418, Edmund Optics). The Köhler lens was mounted on a small linear translational stage (PT1, Thorlabs) to facilitate fine adjustment when focusing the beam at the back focal plane of the microscope objective. Both 3” lenses (the f = 200 mm lens in the telescope and the Köhler lens) were secured on custom 3D printed lens mounts (see supplemental Fig. S1 and supplemental CAD for design).

For Gaussian TIRF/epi-illumination, the galvanometric mirror was set to −20° to send the beam to a periscope mounted on the side of the LS breadboard directing the beam horizontally to a height compatible with alignment on the optical table. The Gaussian beam was then minified and collimated by a lens telescope (f = 150 mm, LA1417-A and f = 25 mm, LA1951-A, both Thorlabs). A second periscope and telescope (f = 75 mm, AC508-075-A and f = 150 mm, AC508-150-A, both Thorlabs) were used for final adjustment of beam height and size. A removable mirror on a magnetic mount was placed on the FF translational stage and was used to direct the beam toward the Köhler lens, which was shared between the Gaussian and FF paths.

The fluorescence emitted from the sample was spectrally filtered (ZT405/488/561/640rpcV3 3 mm thick dichroic mirror in a Chroma BX3 cube; notch filters: ZET642NF, ZET561NF; all from Chroma), collected by the microscope objective lens, and focused by the microscope tube lens to an intermediate image plane from which a two-channel 4f system was aligned in order to access the Fourier plane of the emission path for PSF engineering. The first lens (f = 80 mm, AC508-080-A, Thorlabs) of the 4f system was placed one focal length after the intermediate image plane. Next, a dichroic mirror (T660lpxr-UF3, Chroma) was used to transmit far-red light into one path (the “red” channel) and reflect light of shorter wavelengths into a second path (the “green” channel). Transmissive dielectric double helix phase masks (DH-PMs) with 2-μm or 12-μm axial range (Double Helix Optics), were then positioned in the Fourier planes, one focal length from the first 4f lens, for PSF engineering in each channel. The phase masks were mounted on magnetic mounts for easy placement and removal. To facilitate alignment, the magnetic mounts were placed on XYZ translational stages (PT3A, Thorlabs and 460A-XYZ, Newport). The light was then further filtered (green channel: ET605/70m; red channel: ET700/75m; both Chroma) and focused by the second 4f lenses (f = 80 mm, AC508-080-A, Thorlabs) positioned one focal length from the Fourier plane in each channel. The light paths from the two channels were then merged using a D-shaped mirror and imaged on different regions of an sCMOS camera sensor (Orca Fusion BT, Hamamatsu).

## 3. Results

### 3.1 Design and performance of multimodal illumination platform

Our multimodal illumination platform (Fig. 1a) integrates five different illumination modalities – LS, FF and Gaussian TIRF/epi-illumination – to enable the user to conveniently select the most appropriate method according to their target of interest. Fast and easy switching between these modalities is achieved using a galvanometric mirror to direct the beam into either the LS, FF, or Gaussian paths.

To implement LS illumination, we made key improvements to the previously demonstrated TILT3D platform [32]. Our LS is formed using a cylindrical lens, focused by a long working distance illumination objective, and reflected by a mirror into the sample at an 11° downward tilt. In our design the entire LS path, including the laser heads, the cylindrical lens, the illumination objective, and the reflection mirror are mounted horizontally on a robust platform which can be translated in three dimensions, facilitating easy positioning of the LS in the sample (Fig. 1a, see supplemental Fig. S1 and supplemental CAD for full design). This configuration not only reduces vibration and moments in the LS path, but also ensures the complete decoupling of LS movement in the sample plane from any optical elements aligned on the optical table. Alignment in the sample plane is also facilitated in our design because the LS is directed into the sample chamber using a mirror glued to a custom 3D printed mirror mount (see supplemental CAD for design), decoupling LS and sample movements. An 11° downward tilt enables the LS to be introduced into the sample chamber away from the distorting chamber bottom interface, while allowing illumination of entire adherent cells.

The resulting LS using the 647 nm laser has a thickness of 2.7 μm (1/e^2^ beam waist radius), a width of 80 μm (1/e^2^ beam diameter), and a confocal parameter of 48 μm (1/e^2^) (Fig. 1b,c). These dimensions were selected to make the LS compatible with imaging throughout mammalian cells while maintaining good optical sectioning capacity.

To achieve epi- and TIRF illumination with a FF intensity profile, the Gaussian profile of the output laser beam is reshaped using a square-core multimode fiber, along with a mechanical de-speckler to remove speckle introduced by the fiber (Fig. 1a). This approach decouples the movement of the LS breadboard from the FF optical path so that it does not need to be realigned after positioning the LS. Switching between TIRF and epi-illumination is made convenient and robust using a 1D translational stage to position the beam in the back focal plane of the microscope objective lens, which allows switching without the need for realignment of any optics. We integrate a Gaussian illumination beam to the FF translational stage by positioning a removable mirror before the Köhler lens to enable direct comparison between FF and Gaussian modalities in the same sample (Fig. 1d,e). To enable 3D SRI, lens pairs in 4f configuration are positioned on the emission side of the microscope to allow for PSF engineering using transmissive dielectric phase masks for DH-PSFs with 2 μm or 12 μm axial range independently in the green and red color channels (Fig. 1f). The DH-PSF consists of two lobes instead of one, where the midpoint between the lobes codes for the XY position and the angle between the lobes codes for the Z position. The 2 μm axial range DH-PSFs were used for single-molecule detection in both channels and for fiducial bead detection when imaging close to the coverslip. The 12 μm axial range DH-PSF was used for fiducial bead detection in the green channel during whole-cell single-molecule detection in the red channel.

### 3.2 Light sheet and flat-field TIRF illumination improve the contrast for cellular imaging

The setup performance was benchmarked in terms of contrast improvement by comparing the different illumination modalities for cellular imaging. Conventional Gaussian and FF illumination were used in both epi- and TIRF configurations for imaging of paxillin immunolabeled with CF568 in U-2 OS cells (Fig. 2a, see supplemental Fig. S2 for images with normalized contrast). Paxillin is a focal adhesion protein which is concentrated near the substrate and is thus a suitable target for TIRF illumination. LS illumination was compared to epi-illumination in U-2 OS cells labeled for f-actin using phalloidin conjugated with AF647 (Fig. 2b). Actin is found throughout the cell and can contribute to high out-of-focus background fluorescence.

**Fig. 2.**
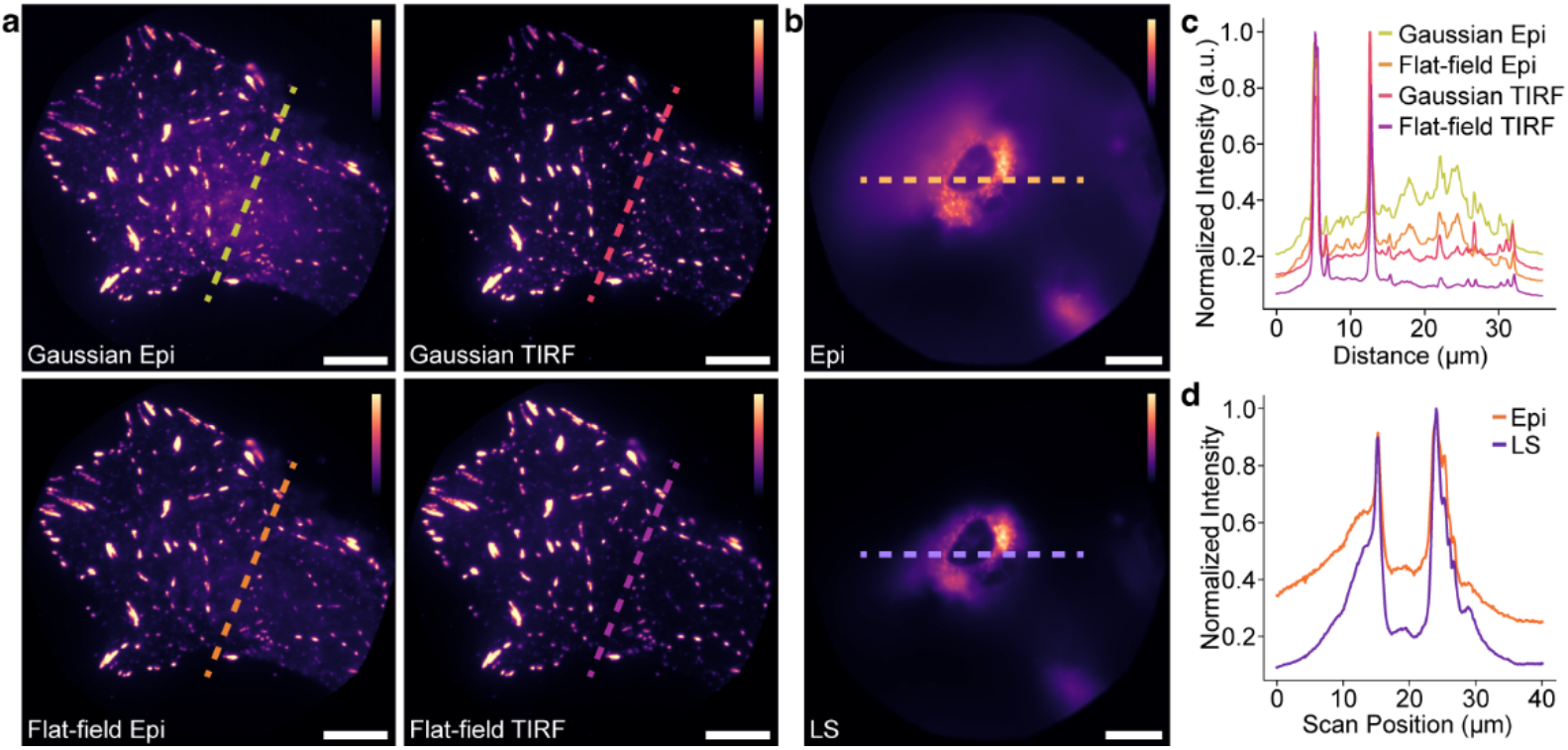
Light sheet (LS) and flat-field (FF) TIRF illumination improve the contrast for cellular imaging. (a) Representative images of a U-2 OS cell immunolabeled with CF568 for paxillin and illuminated using Gaussian epi-illumination (top left), Gaussian TIRF illumination (top right), flat-field (FF) epi-illumination (bottom left), and FF TIRF illumination (bottom right). (b) Representative images of a U-2 OS cell where actin is labeled using phalloidin conjugated with AF647 and illuminated using epi-illumination (top) and light sheet (LS) illumination (bottom). (c) Graph showing line scans as indicated by the dashed lines in (a), demonstrating the contrast improvement and the more homogeneous illumination achieved when using FF TIRF illumination compared to the other illumination modalities. (d) Graph showing lines scans as indicated by the dashed lines in (b), demonstrating the significant contrast improvement when using LS illumination compared to epi-illumination. All scale bars are 10 μm. Color bars indicate the linear color scale used. Each image’s contrast is normalized independently.

For the comparison between Gaussian and FF epi- and TIRF illumination, representative line scans show a clear trend in signal-to-background ratio, where Gaussian epi-illumination results in the lowest ratio and FF TIRF results in the highest – an improvement of up to 4X compared to Gaussian epi-illumination (Fig. 2c). The two TIRF modalities yield, as expected, better contrast than the two epi-illumination modalities. The more homogeneous illumination can also be clearly seen when comparing the two FF modalities to the two Gaussian modalities, where the Gaussian illumination profiles result in higher intensities of fluorescently-tagged paxillin located in the center of the field of view compared to paxillin located at the outer parts of the field of view, whereas the FF modalities result in uniform intensities throughout the field of view.

For the LS comparison to epi-illumination, a representative line scan acquired when imaging actin in the middle of a cell show a signal-to-background improvement of more than 2X when using LS illumination compared to epi-illumination (Fig. 2d).

### 3.3 Flat-field TIRF illumination provides more uniform performance across the field of view for single-molecule localization microscopy

To quantify the improvement in single-molecule localization performance using FF compared to Gaussian illumination, CF568 fluorophores were spin-coated onto a coverslip, imaged in a reducing and oxygen-scavenging buffer using the two illumination modalities, and then localized using ThunderSTORM. The localizations were then quantified in terms of their localization uncertainty (precision), number of localizations, and intensity (signal photons per localization) across the field of view under FF and Gaussian TIRF illumination. Measurements were repeated three times for each modality, and each data set was split into six annuli of equal area for comparison of the statistics in different parts of the field of view (Fig. 3a,b).

**Fig. 3.**
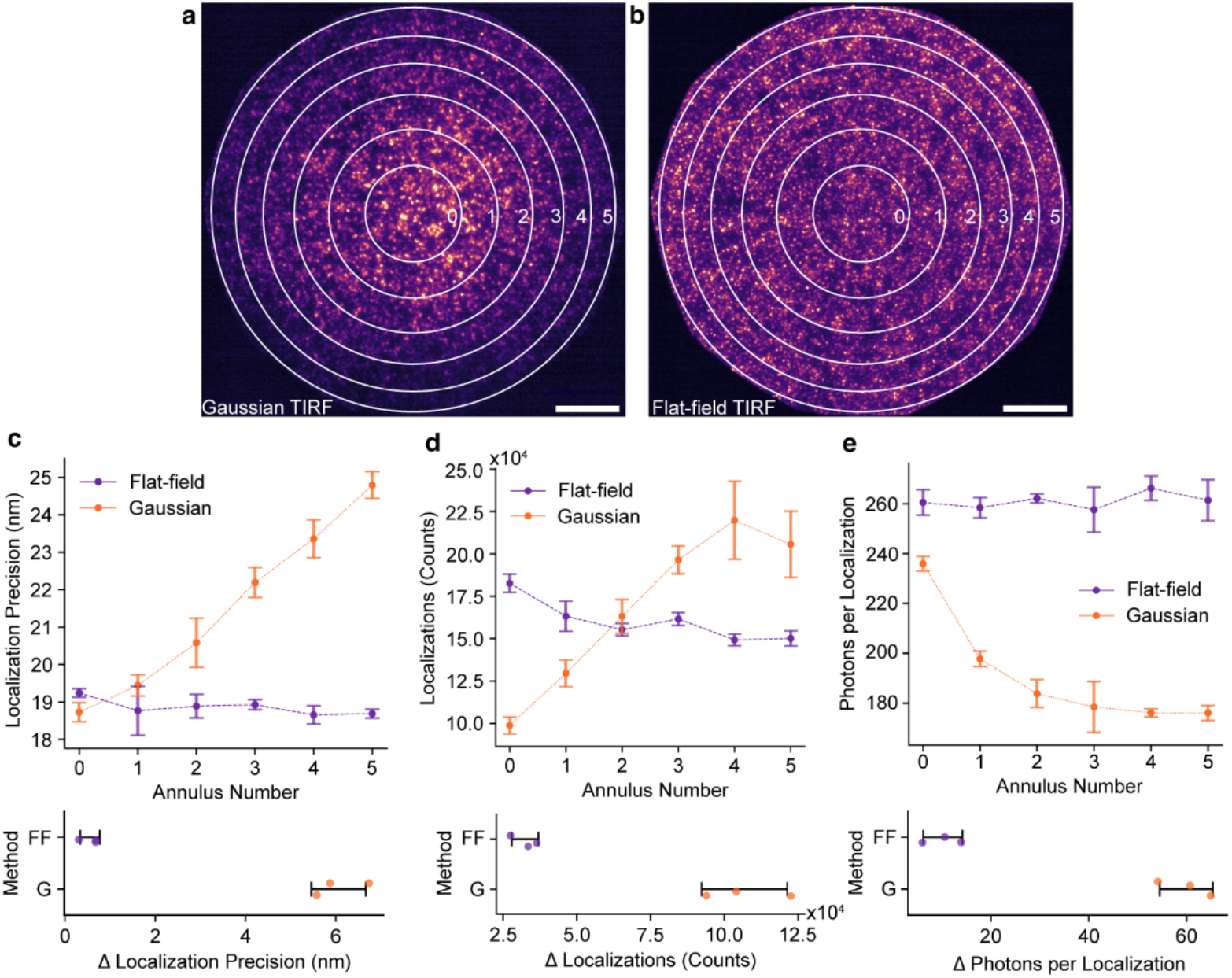
Flat-field TIRF illumination provides more uniform performance across the field of view for single-molecule localization microscopy. (a,b) A dense layer of CF568 fluorophores imaged with (a) Gaussian TIRF illumination and (b) FF TIRF illumination showing the field of view split into six annuli of equal area for parameter quantification. Scale bars are 10 μm. (c-e) Graphs showing Gaussian and FF TIRF comparison across annuli (top panel) and overall difference between annulus 6 and 0 (bottom panel) of the (c) localization uncertainty, (d) number of localizations, and (e) the number of photons per localization (intensity) from single-molecule imaging. The error bars show mean ± standard deviation from three measurements for each illumination modality.

FF illumination results in consistent localization uncertainty (Fig. 3c), number of localizations (Fig. 3d), and localization intensity (Fig. 3e) throughout the field of view, while Gaussian illumination exhibits higher localization uncertainty, more localizations, and fewer photons per localization toward the edge of the field of view compared to the center, due to heterogeneous illumination and photobleaching from the center to the edge. Quantification of the difference, Δ, in the parameters between the outermost and innermost annuli (annuli 6 and 0, respectively), resulted in values for FF compared to Gaussian illumination of 0.6 ± 0.2 nm and 6.1 ± 0.6 nm for the localization precision, 32779 ± 4535 and 106859 ± 14531 for the total number of localizations, and 10.1 ± 4.0 photons and 59.9 ± 5.4 photons for the signal photons per localization, respectively, reported as mean ± standard deviation for three measurements per illumination modality. The homogeneous illumination offered by FF illumination, resulting in consistent SMLM statistics across the field of view, simplifies quantitative analysis and conclusions from SMLM data sets.

### 3.4 3D single-molecule super-resolution imaging throughout mammalian cells

The 3D SRI capabilities of our setup were demonstrated by two-color dSTORM imaging of U-2 OS cells labeled for f-actin and paxillin, where paxillin functions in part as a signal mediator between the extracellular matrix and the actin cytoskeleton [43].

First, actin immunolabeled with AF647 was imaged using LS illumination for optical sectioning throughout the cell. Multiple overlapping slices were illuminated and imaged as needed to cover the cell, where single-molecule data was acquired in the red channel using the 2 μm axial range DH-PSF and the fiducial bead at the coverslip was detected in all slices using the 12 μm axial range DH-PSF in the green channel. After every 20 frames during acquisition, the fiducial bead was illuminated and imaged for one frame using 560 nm FF TIRF illumination. Detection of the fiducial bead with the long-range PSF facilitated both 3D drift correction and easy stitching of slices in post-processing. Next, paxillin, labeled with phalloidin conjugated with CF568, was imaged in the green channel using FF TIRF illumination for optimized contrast at the coverslip, using the same 2 μm DH-PSF for detection of both the single-molecule data and the fiducial bead. These actin and paxillin data sets were then analyzed and stitched together into a single 3D reconstruction (Fig. 4a-d). For whole-cell actin imaging using AF647 and LS illumination, the median XY and Z localization precision was 8.7 nm and 13.1 nm, respectively (supplemental Fig. S3). For paxillin imaging using CF568 and FF TIRF illumination, the median XY and Z localization precision was 9.4 nm and 14.0 nm, respectively (supplemental Fig. S3).

**Fig. 4.**
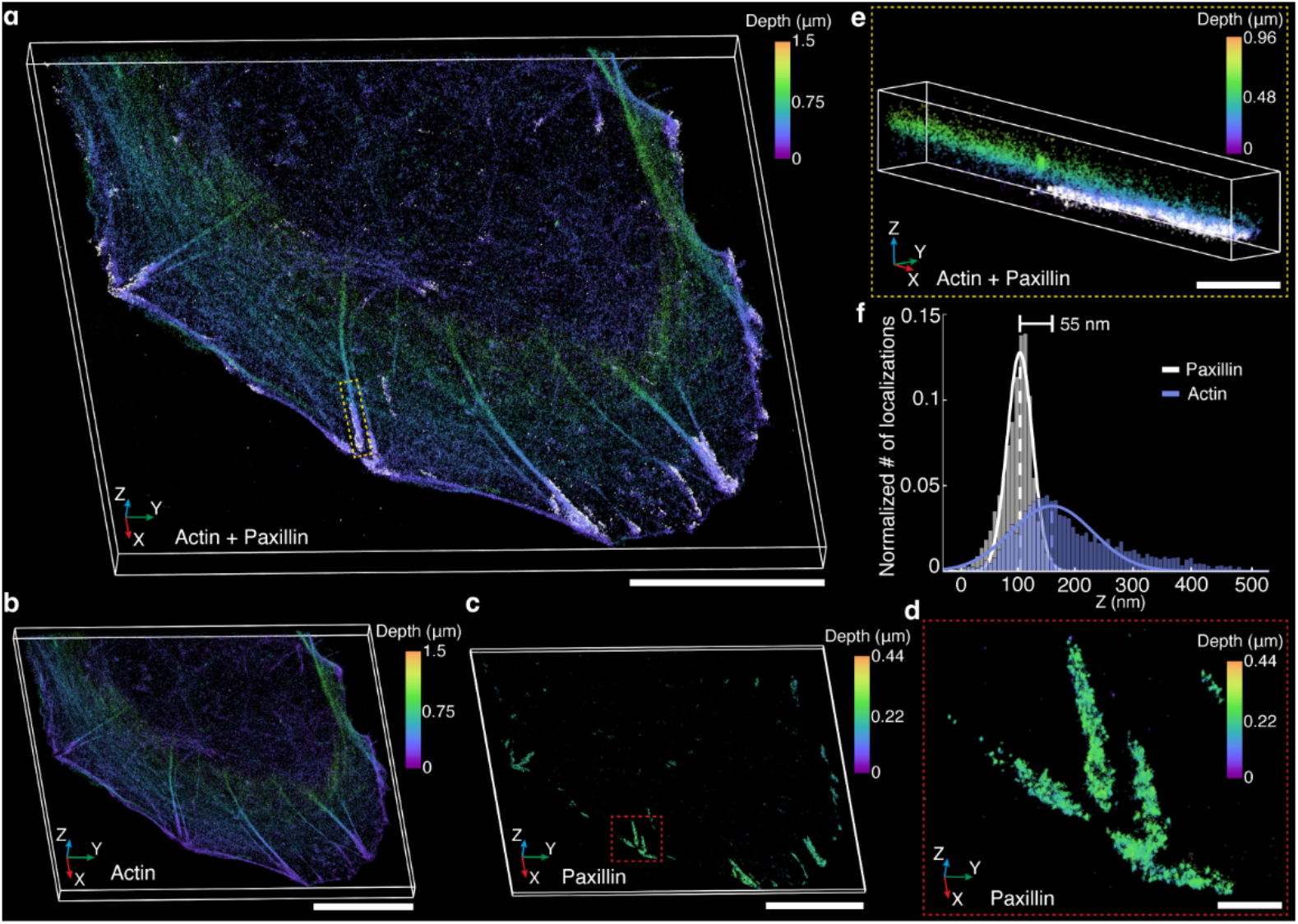
3D single-molecule super-resolution imaging throughout mammalian cells. (a) 3D super-resolved reconstructions of actin (color coded based on Z position) and paxillin (white) in U-2 OS cell. (b) 3D super-resolved reconstruction of the actin shown in (a). (c) 3D super-resolved reconstruction of the paxillin shown in (a), here color coded by depth. (d) Zoom in of the paxillin shown in the red dashed box in (c). (e) Zoom in on the actin and paxillin connection shown in the yellow dashed box in (a). (f) Histogram of Z positions of actin and paxillin localizations of the representative example shown in (e) and corresponding Gaussian fittings used to determine the separation between the two targets. Scale bars are 10 μm in (a), (b), and (c), and 1 μm in (d) and (e).

Next, the separation between actin filaments and paxillin in the focal adhesion complexes was quantified (Fig. 4e,f), which yielded distances of 54.9 ± 6.4 nm (n = 10) (see supplemental Fig. S4 for histograms of the Z distributions of all analyzed actin and paxillin connecting regions), in close agreement with previously reported values [44,45], demonstrating the quantitative performance of this setup for 3D imaging at the nanoscale.

## 4. Discussion

Our multimodal illumination platform integrates fast and simple switching between five different illumination modalities: LS, and FF/Gaussian TIRF and epi-illumination. The modularity in the platform design is achieved by decoupling the LS illumination from the FF modalities using a square-core multimode fiber. In addition, the use of a large square-core multimode fiber simplifies the alignment of the beam into the FF path. Furthermore, the square-core fiber has superior speckling properties compared to circular-core fibers, resulting in uniform illumination [46]. Mounting the optical elements necessary for TIRF and epi-illumination on a 1D translational stage, both for FF and Gaussian modalities, facilitates fast and robust switching between these illumination methods. The combination of the elevated breadboard and the custom 3D printed mirror mount greatly facilitates LS alignment in the sample plane in our implementation.

The different illumination modalities combined with the two-channel 4f system for PSF engineering makes possible the optimization of imaging conditions for studies of a wide range of biological targets. Both modalities of epi-illumination in our setup make fluorophore conversion to the dark state possible throughout the sample, facilitating dSTORM imaging. We demonstrate that for cellular targets in close proximity to the coverslip, FF TIRF illumination enhances overall single-molecule data quality by combining optical sectioning with homogenization of localization uncertainty, localization counts, and the number of photons per localization. LS illumination is then an excellent alternative for optical sectioning of targets that extend throughout the cell above the coverslip.

This multimodal illumination platform is based on primarily commercially available parts, and CAD files are available for the setup and for all custom parts to facilitate easy implementation. We here demonstrate the performance for dSTORM imaging, but the platform is compatible and can easily be implemented for other SMLM modalities, such as (fluorescence) Photoactivated Localization Microscopy ((f) PALM) [2,4] and DNA Points Accumulation in Nanoscale Topography (PAINT) [47]. The platform can also be combined with stage-top incubators, making it compatible for live-cell single-molecule tracking. We think the versatility of our platform will make it a valuable tool for a wide range of future SMLM applications, including quantitative SRI and single-molecule tracking.

## Acknowledgments

We thank Ivana Hsyung for help with cell culturing and labeling and Gabriella Gagliano and Nahima Saliba for helpful discussions. We acknowledge help from Eliberto Batres and Atilla Thomazy with fabrication of custom stage inserts.

## Data availability

Data presented in this paper can be obtained from the authors upon reasonable request.

## Code availability

Calibration and fitting analysis of 2 μm and 12 μm range DH-PSF images were analyzed using a modified version of the open-source Easy-DHPSF software [48,49] (https://doi.org/10.21203/rs.2.9151/v2). Single-molecule data for blinking statistics was analyzed using the open-source ImageJ plugin ThunderSTORM [50] (https://github.com/zitmen/thunderstorm/releases/tag/v1.3/). The custom codes for automatic control of shutters and galvanometric mirror, area-based analysis of blinking data, drift-correction, red-to-green channel transformation, and the modified versions of Easy-DHPSF compatible with 2 μm and 12 μm axial range DH-PSFs are available upon request.

## Supplemental document

See supplemental document for supplemental figures and supplemental CAD files for designs associated with this work.

## References

1. A. Sharonov and R. M. Hochstrasser, “Wide-field subdiffraction imaging by accumulated binding of diffusing probes,” Proc. Natl. Acad. Sci. USA 103, 18911–18916 (2006).

2. E. Betzig, G. H. Patterson, R. Sougrat, O. W. Lindwasser, S. Olenych, J. S. Bonifacino, M. W. Davidson, J. Lippincott-Schwartz, and H. F. Hess, “Imaging Intracellular Fluorescent Proteins at Nanometer Resolution,” Science 313, 1642–1645 (2006).

3. M. J. Rust, M. Bates, and X. Zhuang, “Sub-diffraction-limit imaging by stochastic optical reconstruction microscopy (STORM),” Nat Methods 3, 793–796 (2006).

4. S. T. Hess, T. P. K. Girirajan, and M. D. Mason, “Ultra-High Resolution Imaging by Fluorescence Photoactivation Localization Microscopy,” Biophysical Journal 91, 4258–4272 (2006).

5. L. E. Weiss, J. F. Love, J. Yoon, C. J. Comerci, L. Milenkovic, T. Kanie, P. K. Jackson, T. Stearns, and A.-K. Gustavsson, “Chapter 4 - Single-molecule imaging in the primary cilium,” in Methods Cell Biol, J. M. Bravo-San Pedro and L. Galluzzi, eds. (Academic Press, 2023), Vol. 176, pp. 59–83.

6. L. Möckl and W. E. Moerner, “Super-resolution microscopy with single molecules in biology and beyond– essentials, current trends, and future challenges,” J. Am. Chem. Soc. 142, 17828–17844 (2020).

7. A. M. Sydor, K. J. Czymmek, E. M. Puchner, and V. Mennella, “Super-Resolution Microscopy: From Single Molecules to Supramolecular Assemblies,” Trends in Cell Biology 25, 730–748 (2015).

8. L. von Diezmann, Y. Shechtman, and W. E. Moerner, “Three-Dimensional Localization of Single Molecules for Super-Resolution Imaging and Single-Particle Tracking,” Chem. Rev. 117, 7244–7275 (2017).

9. A.-K. Gustavsson, R. P. Ghosh, P. N. Petrov, J. T. Liphardt, and W. E. Moerner, “Fast and parallel nanoscale three-dimensional tracking of heterogeneous mammalian chromatin dynamics,” MBoC 33, ar47 (2022).

10. T. Kanie, J. F. Love, S. D. Fisher, A.-K. Gustavsson, and P. K. Jackson, “A hierarchical pathway for assembly of the distal appendages that organize primary cilia,” bioRxiv 10.1101/2023.01.06.522944 (2023).

11. H. W. Bennett, A.-K. Gustavsson, C. A. Bayas, P. N. Petrov, N. Mooney, W. E. Moerner, and P. K. Jackson, “Novel fibrillar structure in the inversin compartment of primary cilia revealed by 3D single-molecule superresolution microscopy,” MBoC 31, 619–639 (2020).

12. A. Szymborska, A. de Marco, N. Daigle, V. C. Cordes, J. A. G. Briggs, and J. Ellenberg, “Nuclear Pore Scaffold Structure Analyzed by Super-Resolution Microscopy and Particle Averaging,” Science 341, 655–658 (2013).

13. L. Möckl, K. Pedram, A. R. Roy, V. Krishnan, A.-K. Gustavsson, O. Dorigo, C. R. Bertozzi, and W. E. Moerner, “Quantitative super-resolution microscopy of the mammalian glycocalyx,” Dev. Cell 50, 57–72 (2019).

14. K. Xu, G. Zhong, and X. Zhuang, “Actin, spectrin, and associated proteins form a periodic cytoskeletal structure in axons,” Science 339, 452–456 (2013).

15. B. Huang, S. A. Jones, B. Brandenburg, and X. Zhuang, “Whole-cell 3D STORM reveals interactions between cellular structures with nanometer-scale resolution,” Nat. Methods 5, 1047–1052 (2008).

16. W. E. Moerner and D. P. Fromm, “Methods of single-molecule fluorescence spectroscopy and microscopy,” Review of Scientific Instruments 74, 3597–3619 (2003).

17. A.-K. Gustavsson, P. N. Petrov, and W. E. Moerner, “Light sheet approaches for improved precision in 3D localization-based super-resolution imaging in mammalian cells [Invited],” Opt. Express 26, 13122 (2018).

18. H. D. MacGillavry and C. C. Hoogenraad, “The internal architecture of dendritic spines revealed by super-resolution imaging: What did we learn so far?,” Experimental Cell Research 335, 180–186 (2015).

19. D. Merenich, K. Nakos, T. Pompan, S. J. Donovan, A. Gill, P. Patel, E. T. Spiliotis, and K. A. Myers, “Septins guide noncentrosomal microtubules to promote focal adhesion disassembly in migrating cells,” MBoC 33, ar40 (2022).

20. M. C. Vila, S. Rayavarapu, M. W. Hogarth, J. H. Van Der Meulen, A. Horn, A. Defour, S. Takeda, K. J. Brown,Y. Hathout, K. Nagaraju, and J. K. Jaiswal, “Mitochondria mediate cell membrane repair and contribute to Duchenne muscular dystrophy,” Cell Death Differ 24, 330–342 (2017).

21. S. A. Kamranvar, D. K. Gupta, A. Wasberg, L. Liu, J. Roig, and S. Johansson, “Integrin-Mediated Adhesion Promotes Centrosome Separation in Early Mitosis,” Cells 11, 1360 (2022).

22. M. C. Jones, J. Zha, and M. J. Humphries, “Connections between the cell cycle, cell adhesion and the cytoskeleton,” Phil. Trans. R. Soc. B 374, 20180227 (2019).

23. D. Inoue, D. Obino, J. Pineau, F. Farina, J. Gaillard, C. Guerin, L. Blanchoin, A. Lennon‐Duménil, and M. Théry, “Actin filaments regulate microtubule growth at the centrosome,” The EMBO Journal 38,e99630 (2019).

24. D.-H. Kim, S. B. Khatau, Y. Feng, S. Walcott, S. X. Sun, G. D. Longmore, and D. Wirtz, “Actin cap associated focal adhesions and their distinct role in cellular mechanosensing,” Sci Rep 2, 555 (2012).

25. N. A. Hosny, M. Song, J. T. Connelly, S. Ameer-Beg, M. M. Knight, and A. P. Wheeler, “Super-Resolution Imaging Strategies for Cell Biologists Using a Spinning Disk Microscope,” PLoS ONE 8,e74604 (2013).

26. F. Schueder, J. Lara-Gutiérrez, B. J. Beliveau, S. K. Saka, H. M. Sasaki, J. B. Woehrstein, M. T. Strauss, H. Grabmayr, P. Yin, and R. Jungmann, “Multiplexed 3D super-resolution imaging of whole cells using spinning disk confocal microscopy and DNA-PAINT,” Nat Commun 8, 2090 (2017).

27. C. M. St. Croix, S. H. Shand, and S. C. Watkins, “Confocal microscopy: comparisons, applications, and problems,” BioTechniques 39, S2–S5 (2005).

28. D. Axelrod, “Total Internal REFlection Fluorescence Microscopy in Cell Biology,” Traffic 2, 764–774 (2001).

29. G. Gagliano, T. Nelson, N. Saliba, S. Vargas-Hernández, and A.-K. Gustavsson, “Light Sheet Illumination for 3D Single-Molecule Super-Resolution Imaging of Neuronal Synapses,” Front. Synaptic Neurosci. 13, 761530 (2021).

30. R. M. Power and J. Huisken, “A guide to light-sheet fluorescence microscopy for multiscale imaging,” Nat Methods 14, 360–373 (2017).

31. J. Huisken, J. Swoger, F. Del Bene, J. Wittbrodt, and E. H. K. Stelzer, “Optical Sectioning Deep Inside Live Embryos by Selective Plane Illumination Microscopy,” Science 305, 1007–1009 (2004).

32. A.-K. Gustavsson, P. N. Petrov, M. Y. Lee, Y. Shechtman, and W. E. Moerner, “3D single-molecule super-resolution microscopy with a tilted light sheet,” Nat Commun 9, 123 (2018).

33. N. Saliba, G. Gagliano, and A.-K. Gustavsson, “Whole-cell multi-target single-molecule super-resolution imaging in 3D with microfluidics and a single-objective tilted light sheet,” bioRxiv 10.1101/2023.09.27.559876 (2023).

34. A. Mau, K. Friedl, C. Leterrier, N. Bourg, and S. Lévêque-Fort, “Fast widefield scan provides tunable and uniform illumination optimizing super-resolution microscopy on large fields,” Nature Communications 12, 3077 (2021).

35. F. Stehr, J. Stein, F. Schueder, P. Schwille, and R. Jungmann, “Flat-top TIRF illumination boosts DNA-PAINT imaging and quantification,” Nature Communications 10, 1268 (2019).

36. I. Khaw, B. Croop, J. Tang, A. Möhl, U. Fuchs, and K. Y. Han, “Flat-field illumination for quantitative fluorescence imaging,” Opt. Express 26, 15276–15288 (2018).

37. A. Archetti, E. Glushkov, C. Sieben, A. Stroganov, A. Radenovic, and S. Manley, “Waveguide-PAINT offers an open platform for large field-of-view super-resolution imaging,” Nat Commun 10, 1267 (2019).

38. R. Diekmann, Ø. I. Helle, C. I. Øie, P. McCourt, T. R. Huser, M. Schüttpelz, and B. S. Ahluwalia, “Chip-based wide field-of-view nanoscopy,” Nature Photon 11, 322–328 (2017).

39. K. M. Douglass, C. Sieben, A. Archetti, A. Lambert, and S. Manley, “Super-resolution imaging of multiple cells by optimized flat-field epi-illumination,” Nature Photonics 10, 705–708 (2016).

40. J. Y. L. Lam, Y. Wu, E. Dimou, Z. Zhang, M. R. Cheetham, M. Körbel, Z. Xia, D. Klenerman, and J. S. H. Danial, “An economic, square-shaped flat-field illumination module for TIRF-based super-resolution microscopy,” Biophys. Rep. 2, 100044 (2022).

41. K. Kwakwa, A. Savell, T. Davies, I. Munro, S. Parrinello, M. A. Purbhoo, C. Dunsby, M. A. A. Neil, and P. M. W. French, “easySTORM: a robust, lower-cost approach to localisation and TIRF microscopy,” Journal of Biophotonics 9, 948–957 (2016).

42. J. Deschamps, A. Rowald, and J. Ries, “Efficient homogeneous illumination and optical sectioning for quantitative single-molecule localization microscopy,” Opt. Express 24, 28080–28090 (2016).

43. M. C. Brown and C. E. Turner, “Paxillin: Adapting to Change,” Physiological Reviews 84, 1315–1339 (2004).

44. P. Kanchanawong, G. Shtengel, A. M. Pasapera, E. B. Ramko, M. W. Davidson, H. F. Hess, and C. M. Waterman, “Nanoscale architecture of integrin-based cell adhesions,” Nature 468, 580–584 (2010).

45. L. B. Case, M. A. Baird, G. Shtengel, S. L. Campbell, H. F. Hess, M. W. Davidson, and C. M. Waterman, “Molecular mechanism of vinculin activation and nanoscale spatial organization in focal adhesions,” Nat Cell Biol 17, 880–892 (2015).

46. M. C. Velsink, Z. Lyu, P. W. H. Pinkse, and L. V. Amitonova, “Comparison of round-and square-core fibers for sensing, imaging, and spectroscopy,” Opt. Express 29, 6523 (2021).

47. R. Jungmann, M. S. Avendaño, J. B. Woehrstein, M. Dai, W. M. Shih, and P. Yin, “Multiplexed 3D cellular super-resolution imaging with DNA-PAINT and Exchange-PAINT,” Nature Methods 11, 313–318 (2014).

48. M. D. Lew*, A. R. S. Von Diezmann*, and W. E. Moerner, “Easy-DHPSF open-source software for three-dimensional localization of single molecules with precision beyond the optical diffraction limit,” Protocol Exchange (2013).

49. C. Bayas, A. von Diezmann, A.-K. Gustavsson, and W. E. Moerner, “Easy-DHPSF 2.0: open-source software for three-dimensional localization and two-color registration of single molecules with nanoscale accuracy,” (2019).

50. M. Ovesný, P. Křížek, J. Borkovec, Z. Švindrych, and G. M. Hagen, “ThunderSTORM: a comprehensive ImageJ plug-in for PALM and STORM data analysis and super-resolution imaging,” Bioinformatics 30, 2389–2390 (2014).

