## Supplemental Information for "Multimodal illumination platform for 3D single-molecule super-resolution imaging throughout mammalian cells"

### Supplemental Notes

#### *1. Sample preparation for flat-field (FF) profile characterization and blinking statistics*

For FF and Gaussian TIRF profile characterization, fluorescent beads (560/580 nm, F8800, Invitrogen) at 1:10 dilution were added to 1% polyvinyl alcohol (#17951, Polysciences, Inc.) in nanopure water. 75 μL of this solution was then spin-coated (WS-650, Laurell Technologies) onto high precision coverslips (474030-9020-000, #1.5H, Carl Zeiss Microscopy).

For FF and Gaussian blinking statistics characterization, 1 μL of antibodies conjugated to Alexa Fluor 647 (AF647) fluorophores (ab190573, Abcam) were diluted in 99 μL of reducing and oxygen-scavenging buffer [1]. Briefly, to prepare the buffer, 14 mg glucose oxidase (Sigma Aldrich, G2133) and 50 μL catalase suspension (Millipore Sigma, C100) were added to 200 μL 1X PBS (SH3025601, Cytiva). This solution, (henceforth referred to as "GLOX") was centrifuged for 5 min at 1000 x g and 4ºC and the pellet was discarded. Finally, 10 μL of GLOX was combined with 670 μL nanopure water, 200 μL 50% (w/v) glucose (50-99-7, Sigma Aldrich), 100 μL 1M Tris-HCl (J22638.K2, ThermoFisher Scientific), and 20 μL β-mercaptoethanol (60-24-2, Sigma Aldrich). Next, 7.5 μL of the complete fluorophore-buffer solution was added to 67.5 μL of polyvinyl alcohol, which was then spin-coated onto a high-precision coverslip (474030-9020-000, #1.5H, Carl Zeiss Microscopy) that had been cleaned with compressed nitrogen to remove dust and then plasma-etched (PDC-32G, Harrick Plasma) with atmospheric oxygen for 15 min before the solution was added.

#### *2. Sample preparation for characterization of light sheet (LS) dimensions*

A high-precision coverslip (474030-9020-000, #1.5H, Carl Zeiss Microscopy) was cleaned with compressed nitrogen to remove dust and then plasma-etched (PDC-32G, Harrick Plasma) with atmospheric oxygen for 15 min. A four-sided glass cuvette (SC-4C4S-12.5WG, Mizar Imaging) was then bonded to the coverslip with a small amount of silicone rubber (84431, Smooth-On, Inc.). The fluorescent solution imaged in the LS chamber was prepared by adding phalloidin conjugated to AF647 (A22287, ThermoFisher) to nanopure water at a 91.2 nM concentration.

#### *3. Cell culture and seeding*

Human osteosarcoma cells (U-2 OS, ATCC) were cultured in flasks (FB012935, Fisher Scientific) containing filtered Dulbecco's Modified Eagle Medium (DMEM, 21-063-029, Fisher Scientific), 1 mM sodium pyruvate (11-360-070, Fisher Scientific), and 10% fetal bovine serum (16-000-044, Fisher Scientific) at 37°C and 5% CO2. Cells were plated on high-precision coverslips (474030-9020-000, #1.5H, Carl Zeiss Microscopy) in 6-well plates (08-772-1G, Fisher Scientific) 3-4 days before the experiment. Prior to plating, coverslips were sonicated in acetone for 30 min, dried with compressed nitrogen, and plasma-etched with argon for 15 min.

#### 4. Cell fixation and labeling

U-2 OS cells were washed twice with 1X PBS (SH3025601, Cytiva) and incubated for 20 min in chilled 4% (w/v) formaldehyde solution, made by dissolving paraformaldehyde (15710, Electron Microscopy Sciences) in 1X PBS. The cells were then washed with 1X PBS and incubated at room temperature (RT) for 10 min with 10 mM ammonium chloride (213330, Sigma-Aldrich).

For diffraction-limited imaging of actin, fixed U-2 OS cells were first washed three times with 1X PBS prior to being permeabilized with 0.05% Triton X-100 (X100, Sigma-Aldrich) in 1X PBS for 15 minutes. The cells were then incubated at RT for 20 min in 66 nM phalloidin conjugated with AF647 (A22287, ThermoFisher) in 1X PBS. After incubation, the cells were washed three times with 1X PBS.

For diffraction-limited imaging of paxillin and for two-target SRI of paxillin and actin, fixed U-2 OS cells were first permeabilized by washing three times with 0.2% (v/v) Triton X-100 in 1X PBS with a RT incubation period of 5 min between each wash. Blocking was then performed using 3% (w/v) bovine serum albumin (A2058, Sigma-Aldrich) in 1X PBS for 1 h. The cells were labeled for paxillin with 1:1000 mouse monoclonal primary antibodies (sc-365379, Santa Cruz Biotechnology) in 1% (w/v) bovine serum albumin in 1X PBS overnight at 4°C. Next, the cells were washed three times with 0.1% Triton X-100 with a 3 min incubation between washes. A 1:1000 solution of donkey anti-mouse IgG (H+L) secondary antibodies conjugated with CF568 (20105, Biotium) in 1% (w/v) bovine serum albumin in PBS was added and the cells were left to incubate at RT for 1 h. The cells were washed five times with 0.1% Triton X-100. For two-target SRI of paxillin and actin, the cells were then labeled for actin after the paxillin labeling was completed. The cells were incubated at RT for 20 min in 13.2 nM phalloidin conjugated with AF647 (A22287, ThermoFisher) in 1X PBS. Finally, the cells were washed ten times with 1X PBS.

#### 5. Camera specifications and calibration

The effective pixel size of our system was calibrated by imaging of a 600 line pairs per mm Ronchi ruling (38-566 600 lp/mm, Edmund Optics) using the sCMOS camera (ORCA Fusion BT, Hamamatsu) which resulted in a calibrated pixel size of 128.8 nm in the green channel and 129.3 nm in the red channel. The average conversion gain was calculated as 0.24 photoelectrons per analog-to-digital (A/D) count, and the base level was calculated as 401.2 A/D counts using the ACCéNT sCMOS calibration tool [2]. For all data sets, unless stated otherwise, the camera readout mode was set to “Standard”, binning was set to 2x2, and hot pixel correction mode was set to “Normal.” HCImageLive (Hamamatsu) and Micromanager [3,4] were used interchangeably to control these camera settings and the exposure time.

#### 6. Imaging settings for quantifying the LS and FF characteristics and blinking statistics

To quantify the LS dimensions, images were acquired using a 20X objective (1-U2B225, Olympus) and a CMOS camera (CS235MU, Thorlabs). Calibration of the pixel size of the CMOS camera using the 20X magnification objective was performed using a 600 line pairs per mm Ronchi ruling. Stacks of 100 frames were acquired for each LS orientation (side and top view) at 10 ms exposure time and 8 dB camera gain using the 647 nm laser at 0.9 W/cm$^2$. The same laser wavelength and intensity were used in diffraction-limited imaging of actin in cells illuminated with LS, whereas for epi-illumination the intensity was 1.25 W/cm$^2$. In both cases, the cells were imaged with the sCMOS camera with an exposure time of 50 ms.

The data for the fluorophore blinking statistics comparison between FF and Gaussian illumination was acquired using the sCMOS camera "Fast" readout mode and an exposure time of 5 ms to balance temporal resolution with single-molecule localization precision. Multiple 4000-frame stacks of images were acquired at multiple fields of view using the 560 nm laser at roughly matched illumination intensities of 250 mW/cm$^2$. For each field of view, acquisition began as soon as the shutter opened, allowing us to capture initial shelving and subsequent photobleaching. The IX-83 microscope automatic focus locking functionality was used for each of these data sets to ensure consistent focus between fields of view. The same laser intensities were used for imaging the dense layer of fluorescent beads and for diffraction-limited imaging of paxillin in cells when using TIRF illumination, whereas 150 mW/cm$^2$ was used for epi-illumination. For this imaging, the sCMOS camera exposure time was set to 50 ms and "Standard" readout mode.

#### *7. Imaging settings for two-color 3D single-molecule super-resolution microscopy*

For two-color 3D SRI of actin and paxillin in U-2 OS cells, the sample was placed on the microscope and 3 μL of fiducial bead solution (560/580 nm, 0.2 μm, F8800, Invitrogen, diluted 1:10000 in PBS) were added dropwise until a suitable number of beads was visible in a given field of view while the sample was monitored using the 560 nm laser at 3.4 W/cm$^2$. Prior to beginning SRI, the PBS buffer was exchanged with the reducing and oxygen-scavenging buffer detailed in Supplemental Note 1 to facilitate direct Stochastic Optical Reconstruction Microscopy (dSTORM) [5] imaging.

For 3D SRI of actin, a DH-PM with 2 μm axial range (DH1-670-3249, Double Helix Optics) was placed in the red path of the 4f system and a DH-PM with 12 μm axial range (DH12R-580-3249, Double Helix Optics) was placed in the green 4f path for fiducial bead detection. Single-molecule data was acquired with LS illumination using the 647 nm laser at 3.8 kW/cm$^2$. The fiducial bead was imaged with the 560 nm laser at 3.4 W/cm$^2$ using FF epi-illumination every 20 frames. The laser shutters and the galvanometric mirror were controlled through the multifunction I/O device using a custom MATLAB script to enable fast, automated switching between LS illumination at 647 nm and FF epi-illumination at 560 nm.

After completing the data acquisition for actin, SRI of paxillin was performed with FF TIRF illumination using the 560 nm laser at 390 W/cm$^2$. Since both the single-molecule target and the fiducial bead were excited with the same 560 nm laser and both were located in close proximity to the coverslip, the 12 μm axial range DH-PM was substituted by a 2 μm axial range DH-PM (DH1-580-3240, Double Helix Optics) in the green channel. Prior to the start of single-molecule acquisition, the fiducial bead was imaged for 100 frames using the 12 μm axial range DH-PM in the green channel for later axial registration between targets.

The camera exposure time was set to 50 ms and SRI of both actin and paxillin was performed after converting a large fraction of the fluorophores to a dark state using either FF or Gaussian epi-illumination. For actin and paxillin SRI, the fluorophore conversion to dark state was performed with the 647 nm laser using an intensity of 3.0 kW/cm$^2$ and with the 560 nm laser using an intensity of 160 W/cm$^2$, respectively. For each target, 50000 frames of single-molecule data were acquired.

To perform calibration of the 2 μm and 12 μm axial range double-helix PSFs (DH-PSFs) and registration between the red and green channels, 7.5 μL of fluorescent bead solution (calibration: 560/580 nm: F8800; 660/680 nm: F8807; registration: Tetraspeck T7280; all Invitrogen, diluted 1:10000 in nanopure water) was added to 67.5 μL of 1% polyvinyl alcohol in nanopure water and spun onto a high-precision coverslip (474030-9020-000, #1.5H, Carl Zeiss Microscopy) that had been cleaned with compressed nitrogen to remove dust and then plasma-etched (PDC-32G, Harrick Plasma) with atmospheric oxygen for 15 min before the solution was added. Images were then acquired of the beads while scanning over the 2 μm and 12 μm axial ranges using the piezoelectric XYZ translation stage with 50 nm and 100 nm step size, respectively. For each step 10 frames were acquired. For channel registration, five different fields of view were imaged simultaneously in both channels using a Tetraspeck fiducial bead sample, where 100 frames were acquired per field of view. Additionally, 100 dark frames were acquired for further calibration and registration analysis.

#### *8. Image analysis for quantification of LS and FF characteristics and blinking statistics*

To determine the LS dimensions, LS image stacks were averaged using the open-source software ImageJ [6,7]. The image acquired of the focused direction of the LS (the LS thickness) was analyzed using a custom Python script. Line scans perpendicular to the propagation axis were acquired at every row of pixels in the image and fitted with Gaussian functions. The beam waist radius was determined as the $1/e^2$ radius of the Gaussian fit at the LS focus. To determine the confocal parameter of the light sheet, the $1/e^2$ radius of each Gaussian fit was measured, and this sequence of thicknesses along the optical axis was fitted accordingly [8]. The LS width was determined from the $1/e^2$ width of Gaussian fits of the LS perpendicular to the propagation axis from the images acquired of the collimated direction of the LS.

To compare the illumination profile homogeneity across the field of view for the Gaussian and FF TIRF illumination modalities, 100-frame image stacks of beads in each mode were averaged in ImageJ and line scans were acquired for each averaged image.

To compare the illumination profile homogeneity and out-of-focus background between FF and Gaussian TIRF and epi-illumination modalities for cellular imaging, 100-frame image stacks in each mode were averaged in ImageJ and line scans were acquired for each averaged image.

To quantify the contrast improvement between LS and epi-illumination, 100-frame image stacks were acquired of labeled actin in cells in each mode. The stacks were averaged in ImageJ and line scans were acquired for each averaged image.

To compare the blinking statistics throughout the field of view between FF and Gaussian illumination profiles, the open-source ImageJ plugin ThunderSTORM [9] was used for single-molecule localization, where localizations with an uncertainty greater than 100 nm were discarded to facilitate visualization (only around 0.004% of total localizations were discarded for each stack). A custom Python script was used to split each data set into 6 concentric annuli of equal area. The signal photons per localization, the localization precision, and the total number of localized molecules in each annulus were compared for three fields of view each for the two illumination methods.

#### *9. Analysis of 3D single-molecule data*

ImageJ was used to crop the images obtained in both channels for all data. Next, the single-molecule and fiducial bead data were imported into a modified version of Easy-DHPSF [10,11], a MATLAB-based open-source software for DH-PSF single-molecule localization. For the paxillin data, since both the fiducial bead and the single-molecule data were acquired in the same channel, single-molecule localization and drift correction were performed using a modified version of Easy-DHPSF compatible with the 2 μm axial range DH-PSF.

For the actin data, single-molecule localization was performed using the same Easy-DHPSF version as used for paxillin analysis. Since the fiducial bead data for the actin sample was acquired using a 12 μm axial range phase mask in a different spectral channel, the localization of the fiducial bead was performed using a modified version of Easy-DHPSF compatible with 12 μm axial range DH-PSFs. The fiducial bead data was then used to drift correct the actin single-molecule localizations. A custom MATLAB drift correction script was used to fit a cubic smoothing spline function to the XYZ position of the bead over time to correct the drift.

XY registration between the two channels was performed by applying the MATLAB function `imregtform` to the Tetraspeck fiducial bead stacks to obtain an affine matrix transformation that was later applied to the single-molecule localizations obtained from Easy-DHPSF. For Z registration between the two channels, the stack of the paxillin fiducial bead acquired with the 12 μm DH-PSF was localized using the version of Easy-DHPSF compatible with the 12μm range. The Z position of the paxillin fiducial bead was averaged and subtracted from the Z position of the actin fiducial bead from the average of the first 5 frames. All paxillin and actin localizations were scaled by 0.75 [12,13] to account for index mismatch between the glass coverslip and the sample.

After both the paxillin and actin data were localized and drift-corrected, both data sets were stitched together after correcting for the difference in the lateral position of the fiducial bead in the first frame of the paxillin and actin data sets. For rendering and visualization, the 3D single-molecule data was imported into the software Vutara SRX (Bruker), where localizations in each data set were filtered independently for DH interlobe distance and localization precision in Z. Specifically, for the paxillin data, the following filters and settings were applied: 6.9 – 7.9 pixels interlobe distance and 3 nm to 20 nm Z localization precision. For the actin data, the filters and settings were as follows: 7.0 - 9.5 pixels interlobe distance and 3 nm to 25 nm Z localization precision. The filtered localizations were visualized with the point splatting rendering option using a 25 nm particle size. Several regions (n = 10) where actin and paxillin connected were selected in Vutara SRX, and the distributions of localizations in Z were fitted with Gaussian functions. The distance between the actin and paxillin distributions was then quantified as the difference between the peaks of the two Gaussian fits.

### Supplemental figures

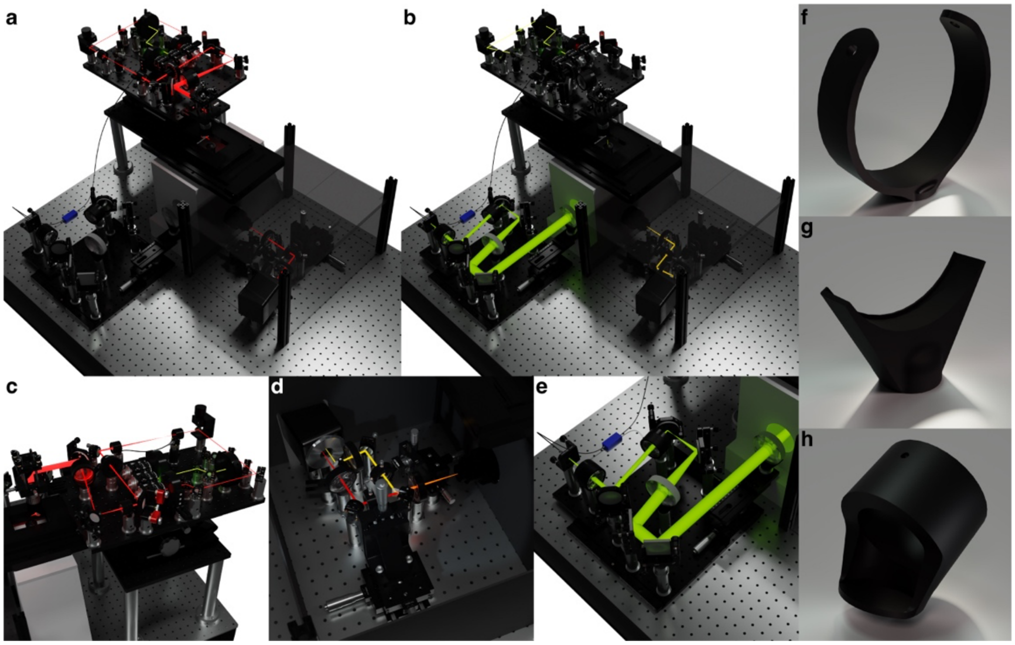


Fig. S1. Selected views of multimodal illumination microscope CAD rendering. (a,b) Rendering of the full setup, where (a) the light sheet (LS) path is illuminated and (b) the flat-field (FF) path is illuminated. In both, the light-proof box surrounding the 4f system path for point spread function (PSF) engineering is rendered as translucent to aid in visualization. (c) The LS breadboard and stages shown in more detail. (d) The 4f system path shown with a schematic representation of how the emission light is split into two channels. (e) The FF TIRF/epi-illumination module of the setup, where both translating stages can be seen, along with the de-speckler (blue). 3D renderings of (f) the custom shutter mount, (g) three-inch lens mount, and (h) 11º LS objective mirror mount.

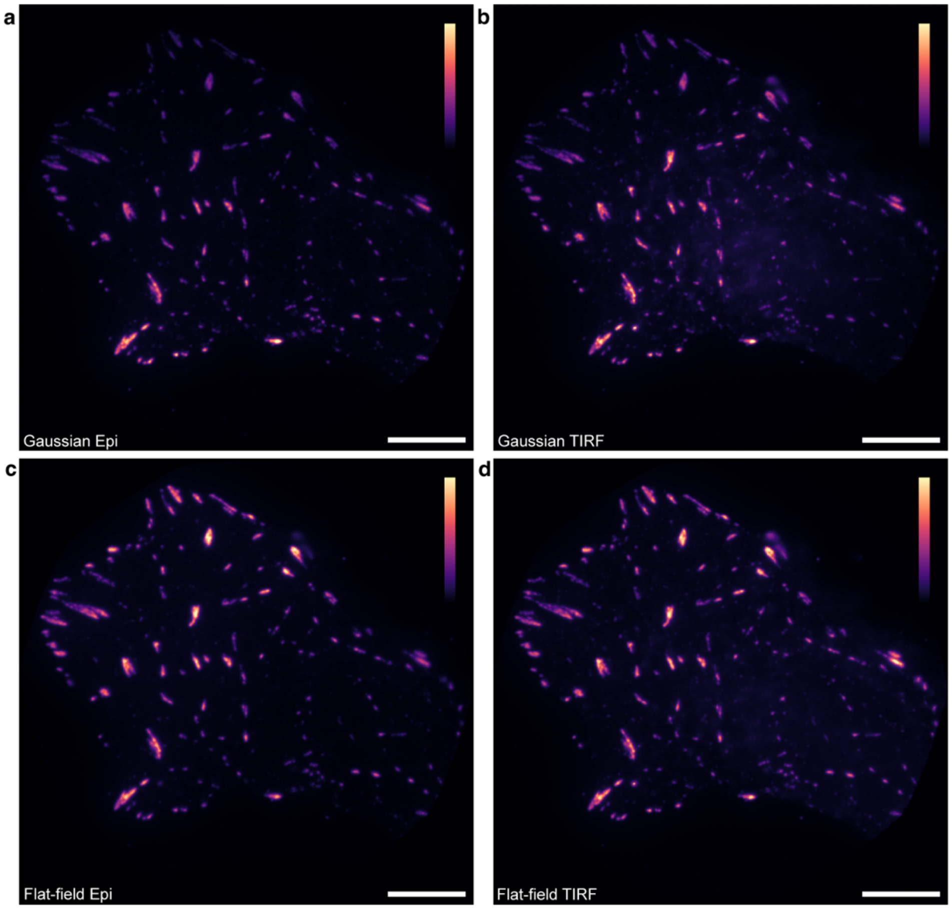


Fig. S2. Diffraction-limited images of paxillin shown in Fig. 2a, here shown with lower contrast, when illuminated with (a) Gaussian epi-illumination, (b) Gaussian TIRF illumination, (c) flat-field epi-illumination, and (d) flat-field TIRF illumination. The contrast in each image is normalized to the highest and lowest values in each respective image. Scale bars are 10 µm. Color bars indicate the linear color scale used.

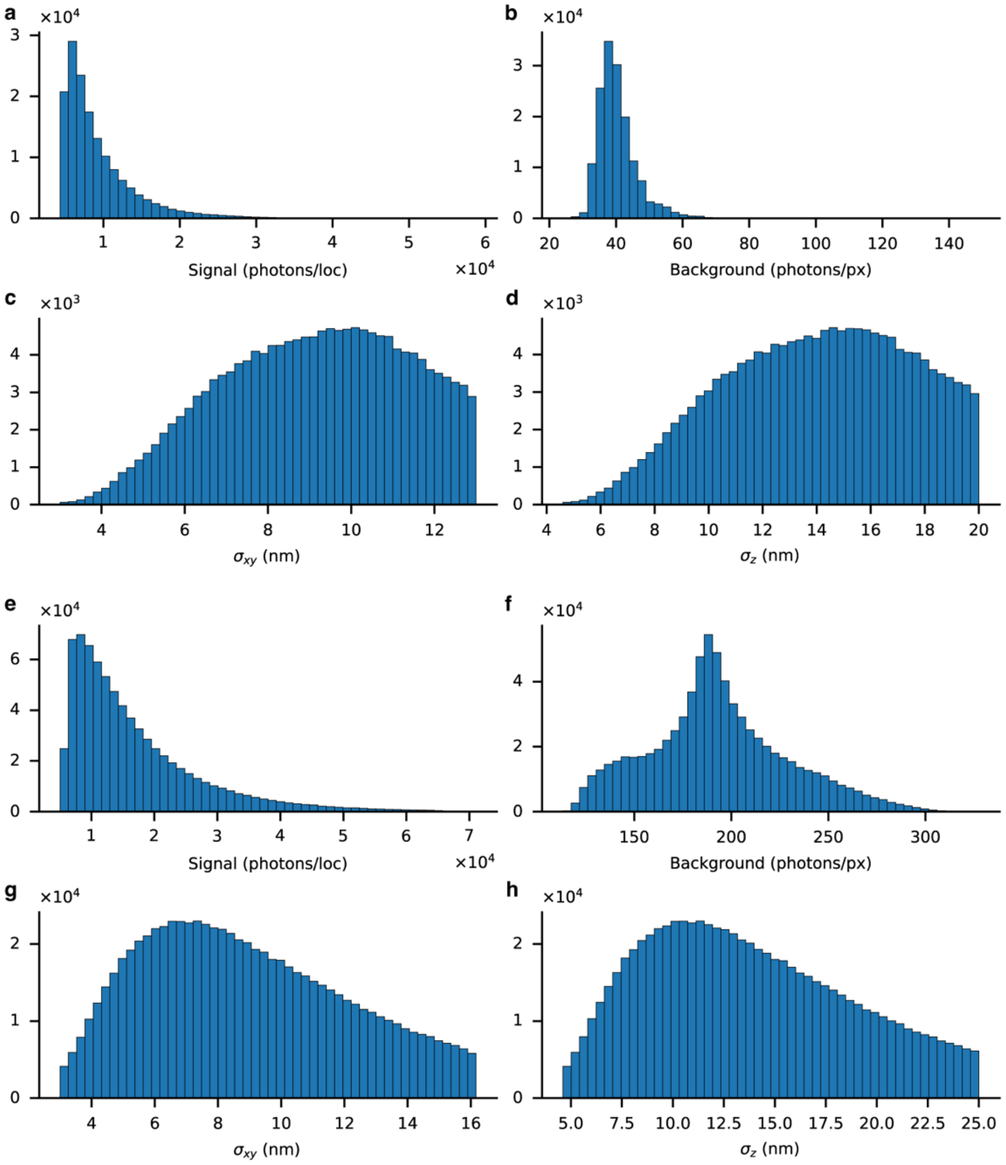


Fig. S3. Histograms showing the (a,e) signal photons per localization, (b,f) background photons per pixel, and (c,g) XY and (d,h) Z localization precision for paxillin (a-d) and actin (e-h) single-molecule data acquired using the double-helix (DH) PSF and shown in Fig. 4. The paxillin data was filtered for localizations with a DH-PSF lobe distance within 6.7 - 7.9 pixels and for Z localization precision between 3 - 20 nm. This resulted in ~ 152000 filtered localizations with median photons per localization of 7765, background photons per pixel of 39, and an XY and Z localization precision of 9.4 nm and 14.2 nm, respectively. The actin data was filtered for DH-PSF lobe distance within 7.0 - 9.5 pixels and for Z localization precision between 3 - 25 nm. This resulted in ~740000 filtered localizations with median photons per localization of 13772, background photons per pixel of 191, and an XY and Z localization precision of 8.7 nm and 13.1 nm, respectively.

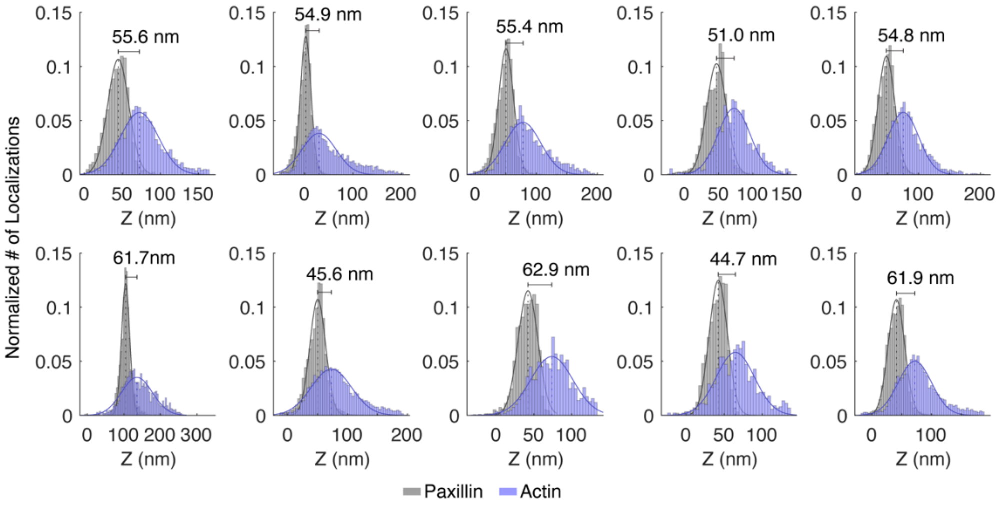


Fig. S4. Histograms of n = 10 connection regions of actin and paxillin. Each histogram shows the z positions of the actin and paxillin localizations and corresponding Gaussian fittings used to determine the separation between the two targets.

### References


1. A. R. Halpern, M. D. Howard, and J. C. Vaughan, "Point by Point: An Introductory Guide to Sample Preparation for Single-Molecule, Super-Resolution Fluorescence Microscopy," Current Protocols in Chemical Biology **7**, 103–120 (2015).
2. R. Diekmann, J. Deschamps, Y. Li, T. Deguchi, A. Tschanz, M. Kahnwald, U. Matti, and J. Ries, "Photon-free (s)CMOS camera characterization for artifact reduction in high- and super-resolution microscopy," Nature Communications **13**, 3362 (2022).
3. A. Edelstein, N. Amodaj, K. Hoover, R. Vale, and N. Stuurman, "Computer Control of Microscopes Using µManager," CP Molecular Biology **92**, (2010).
4. A. D. Edelstein, M. A. Tsuchida, N. Amodaj, H. Pinkard, R. D. Vale, and N. Stuurman, "Advanced methods of microscope control using µManager software," J Biol Methods **1**, e10 (2014).
5. M. Heilemann, S. van de Linde, M. Schüttpelz, R. Kasper, B. Seefeldt, A. Mukherjee, P. Tinnefeld, and M. Sauer, "Subdiffraction-Resolution Fluorescence Imaging with Conventional Fluorescent Probes," Angewandte Chemie International Edition **47**, 6172–6176 (2008).
6. M. D. Abràmoff, P. J. Magalhães, and S. J. Ram, "Image processing with ImageJ," Biophotonics international **11**, 36–42 (2004).
7. C. A. Schneider, W. S. Rasband, and K. W. Eliceiri, "NIH Image to ImageJ: 25 years of image analysis," Nat Methods **9**, 671–675 (2012).
8. A.-K. Gustavsson, P. N. Petrov, M. Y. Lee, Y. Shechtman, and W. E. Moerner, "3D single-molecule super-resolution microscopy with a tilted light sheet," Nat Commun **9**, 123 (2018).
9. M. Ovesný, P. Křížek, J. Borkovec, Z. Švindrych, and G. M. Hagen, "ThunderSTORM: a comprehensive ImageJ plug-in for PALM and STORM data analysis and super-resolution imaging," Bioinformatics **30**, 2389–2390 (2014).
10. M. D. Lew*, A. R. S. Von Diezmann*, and W. E. Moerner, "Easy-DHPSF open-source software for three-dimensional localization of single molecules with precision beyond the optical diffraction limit," Protocol Exchange (2013).
11. C. Bayas, A. von Diezmann, A.-K. Gustavsson, and W. E. Moerner, "Easy-DHPSF 2.0: open-source software for three-dimensional localization and two-color registration of single molecules with nanoscale accuracy," (2019).
12. B. Huang, W. Wang, M. Bates, and X. Zhuang, "Three-Dimensional Super-Resolution Imaging by Stochastic Optical Reconstruction Microscopy," Science **319**, 810–813 (2008).
13. Y. Li, M. Mund, P. Hoess, J. Deschamps, U. Matti, B. Nijmeijer, V. J. Sabinina, J. Ellenberg, I. Schoen, and J. Ries, "Real-time 3D single-molecule localization using experimental point spread functions," Nat Methods **15**, 367–369 (2018).
